# ULTRASTRUCTURAL STUDY OF THE ESOPHAGUS AND STOMACH OF *Arapaima gigas* (Schinz 1822), JUVENILE PAICHE, CREATED EXCAVATED TANK

**DOI:** 10.1101/2021.04.07.438795

**Authors:** Keila Silva Pinto, Luana Félix de Melo, Julia Bastos de Aquino, Jerônimo Vieira Dantas Filho, Maria Angelica Miglino, Sandro de Vargas Schons, Rose Eli Grassi Rici

## Abstract

Paiche (*Arapaima gigas)* belongs to the Kingdom *Animalia*, Phylum *Chordata*, Class *Actinopterygii*, Order *Osteoglossiformes,* Family *Arapaimidae,* Genus *Arapaima,* and its origin may date to the Jurassic period. The species has natural habitat in the Amazonian rivers, found mainly in marginal lakes, being considered an important fishing resource, with high market value and high demand for meat and leather in both Brazilian and international trade. This study aims to describe the morphology of the esophagus and stomach by light microscopy and scanning electronics microscopy. The esophagus was presented as muscular, short, tubular and fan-shaped in the cranial portion, also presenting deep longitudinal folds, and the entire mucosa is covered by mucus secretory cells with distinct morphological characteristics. Pirarurcu’s stomach has a J-shape divided into three regions: cardiac with a lighter aspect, fundus portion with few folds in the mucosa, and pyloric with deeper folds, also presenting gastroliths in fundus and pyloric portions. Both microscopy studies highlighted three glandular regions, composed by mucoid columnar epithelial cells, gastric crypts with different shapes and sizes depending on each portion, in which the different shapes of the mucosal folds in each region of the stomach were evident, and digitiform microsaliences were found in the cardiac region, and micro-orifices and desmosome in the fundus region. Also, fundus and pyloric portions produce more mucus than the cardiac. Then morphology found was consistent with the eating habits and management of distinct characteristics of the digestive tract.

## Introduction

Paiche (*Arapaima gigas* Schinz, 1822) is the largest scale fish of the Amazonian rivers, being found mainly in locations with low or no currents, such as marginal lakes [1]. *A. gigas* is considered the most important and emblematic species of Amazonian ichthyofauna, not only for its large size, but also for the historical and economic context of artisanal fishing [2]. Among the fish of the Order *Osteoglossiformes,* paiche is the best known and, such as others of this order, it is characterized by the presence of parasphenoid bone [3].

However, due to the high exploitation of natural resources and lack of inspection, paiche is in a worrying situation, mainly due to the reduction of natural resources and near disappearance of its habitat [4]. On the other hand, it presents itself as an important fishing economic activity, as it is a neotropical fish, of large size, high rusticity, easy adaptation to high rates of density and high temperatures, as well as good feed conversion and excellent weight gain in low water quality environments [5]. However, knowledge of the fishing biology of these species is necessary, especially, to understand eating peculiarities, aiming to offer rational measures of exploitation [6]. The architecture and the structure of the gastrointestinal tract have been described in several fish species, mainly due to the interest of wide variations in both morphology and functions related to nutritional dynamics, physiological changes in aquaculture target species and digestive pathologies [7–9].

According to Rodrigues and Cargnin-Ferreira [10], the morphological aspects of the digestive tract of the *A. gigas,* in the juvenile phase is similar to other carnivorous teleosts, allowing the species to ingest, store and digest large foods. In particular, such studies are relevant to understand morphophysiological particularities related to seizure, digestion, and absorption of food in target aquaculture species, and thus reduce adaptive flexibility in response to changes in diet, due to the decrease in the variation in nutrient composition in natural environment, administered in captivity production [11].

Based on the aforementioned, the propose of this investigation is to describe the morphological and structural aspects of the esophagus and stomach of the young *Arapaima gigas*, raised in commercial creations in excavated tanks.

## Material and Methods

Thus, esophagus and stomach were collected from six specimens of Paiche (*Arapaima gigas*), in the juvenile phase, raised in an intensive system of tanks excavated and fed with a pelletized commercial diet, from an aquaculture property in the municipality of Pimenta Bueno/Rondônia/Brazil. The fish were caught for the consumption of the property, by artisanal fishing, and after removal of the viscera, the esophagus and stomach were washed in running water, removing the food content present inside and fixed in 10% buffered formaldehyde. The sample was processed at Facilite Advanced Center for Diagnostic Imaging - CADI- FMVZ-USP.

Organ dissection and mucosal exposure were performed on a plank to macroscopically observe the different structures. Then these organs were later documented with a photographic camera. For microscopic analysis, tissue samples were dehydrated in a series of increasing concentrations of ethanol (70 to 100%) and diaphanized in xylol, with subsequent inclusion in histological liquid paraffin and then microtomized at 5μm thick and subsequently stained with Hematoxylin-Eosin (HE). The images were obtained with the Nikon Eclipse E-800 light microscope.

Part of the material was processed for scanning electron microscopy (SEM). Samples fixed in 10% formaldehyde were dehydrated in increasing sets of alcohols at concentrations of 50%, 70%, 90% and 100%, dried in LEICA critical point apparatus in CPD300 (FMVZ-USP), fixed with carbon glue in aluminum metal bases (stub) and metallized (sputting) with gold in the EMITECH K550 metallizer (FMVZ-USP), being analyzed and photographed under a LEO 435VP scanning electron microscope (FMVZ-USP).

The project was presented and approved by the Research Ethics Committee on the use of animals of the Federal University of Rondônia, with protocol No. 043/2019.

## Results

Anatomically, the esophagus and stomach of the *Arapaima gigas* are tubular organs composed of modified muscle fibers that connect the pharyngeal cavity to the proximal intestine (Fig. 1A). The esophagus is presented in form of a muscular and short rectilinear tube (Fig. 1 A, C), but at the pharyngo-esophageal junction (cranial portion), a marked dilution of the fan-shaped organ (a) is observed, and in the caudal portion a visible reduction of the lumen (b). On the other hand, the stomach is a J-shaped muscle sac (Fig. 1 A), located on the left side of the cavity, extending from the esophagus to the proximal bowel portion. The esophageal mucosa is characterized by numerous longitudinal folds, most evident in the medial and caudal region (arrows on Fig. 1 C).

**Fig 1.**
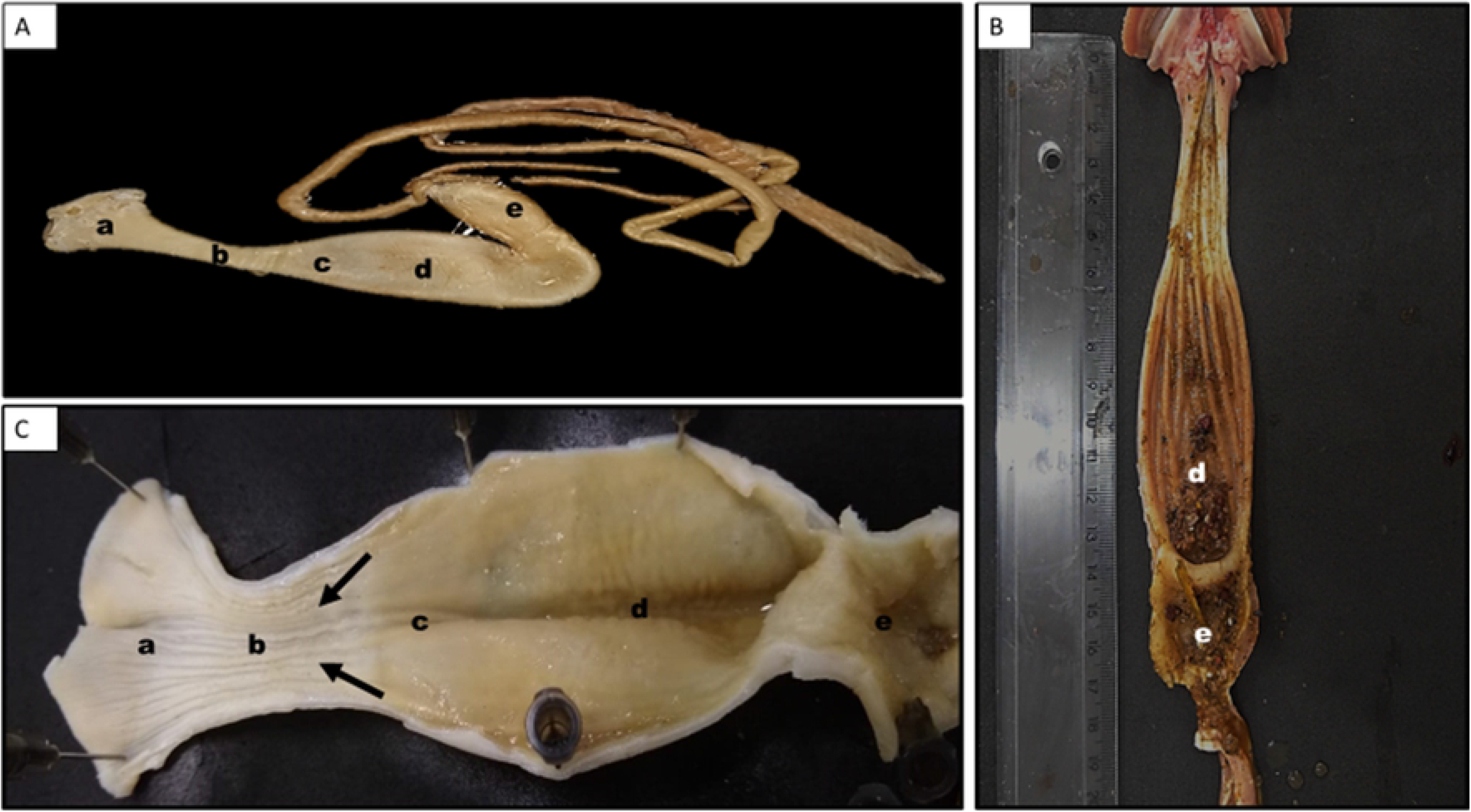
In A, external photomacrography of the *A. gigas* digestive tube, (a) esophageal cranial portion (b) caudal esophageal portion straight, short and muscular tube; (c, d, e) J-shaped stomach. In C mucosa of the esophagus and stomach,(a) fan-shaped esophageal cranial portion (b), medial portion and caudal (arrows) with presence of longitudinal folds in the esophageal mucosa, in (c) cardiac portion of the stomach, lighter aspect, in (d) mucosa in gastric folds of the fundus portion and (e) pyloric portion of the stomach. In B stomach (d, e) fungal and pyloric portions respectively with presence of gastroliths.

As the esophageal mucosa is replaced by gastric, such folds become thicker and more visible. However, after stomach fixation in 10% formaldehyde it was possible to differentiate it into three distinct portions, mainly by the color and shape of the distribution of the gastric folds, in: cranial or cardiac portion, with a lighter aspect and the central sulcus and the mucosa folds parallel to the central axis of the organ (c). In the medial or fundus portion, gastric folds are more evident, with the presence of transverse sulcus (d) and in the caudal or pyloric portion, gastric folds are higher with deep sulcus (e). During the opening of the paiche’s stomach (Fig. 1 B) the presence of small gastroliths was observed in the fungal and pyloric portions (d, e).

In the microscopic study of light, it was observed that the esophagus is wrapped by a stratified epithelium (Fig. 2 A) (line) with three different cell types: the simple epithelial cells, the mucous cells and the claviform cells. The number of layers varied from 1 to 12 layers of cells. The mucous cells presented higher numbers and they were distributed among the layers of the stratified epithelium. By histochemical methods, with hematoxylin and eosin (H/E) two cell types could be differentiated: acidophil and basophil (Fig. 2 A).

**Fig. 2.**
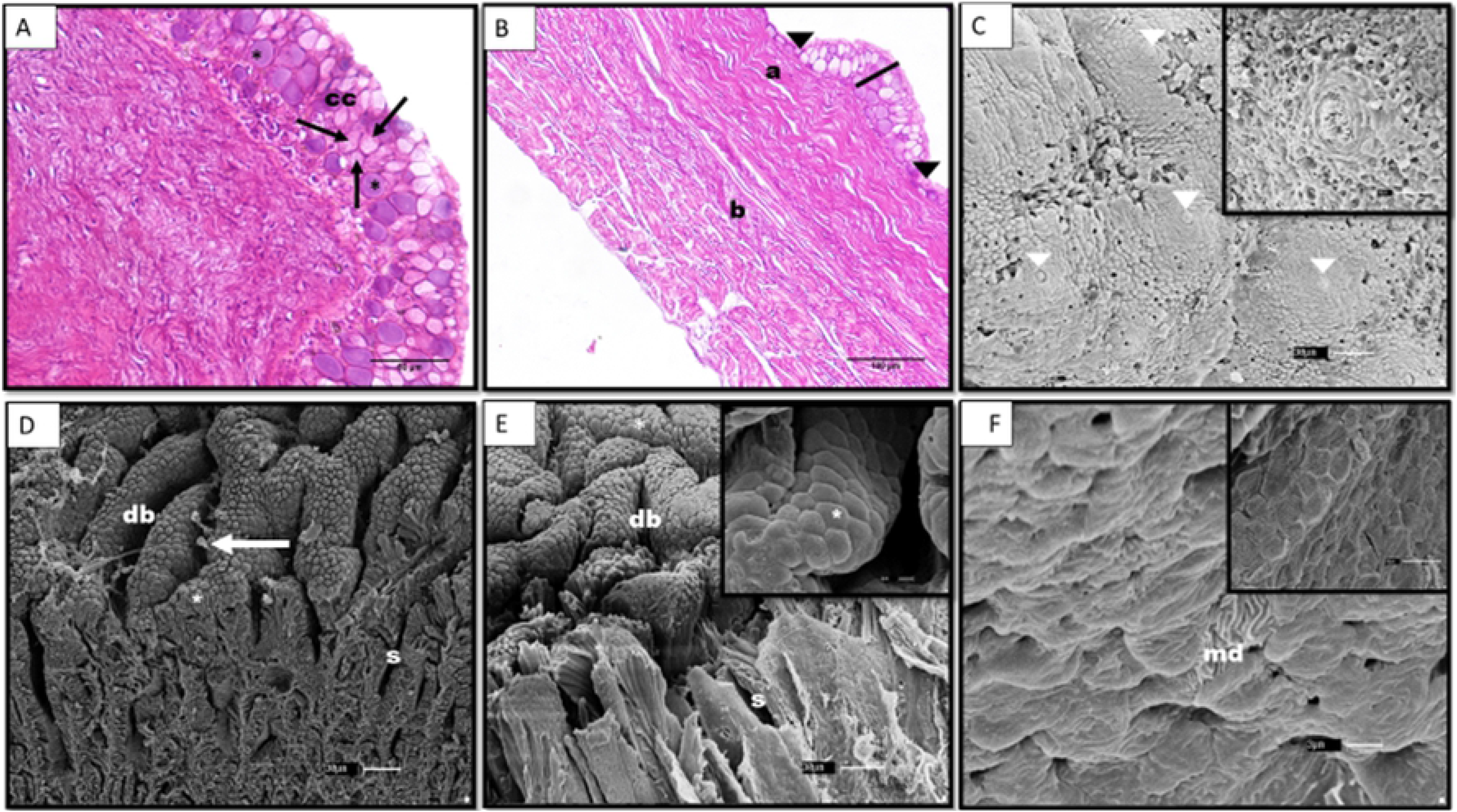
Esophageal photomicrograph *A. gigas*. Cross-section in A, B, (HE)/ML. In A esophageal mucosa lyxiform cells with halo around the nucleus (cc), cuboid cells, granulous aspect (arrows), the major, rounded basophilic cells and basophilic cytoplasm (*). B epithelial cells of prismatic-to cuboids-shape, epithelium is formed by up to two layers of cells (arrowhead), stratified epithelium with the presence of secretory epithelial cells (trace), mucosal and submucosa lamina propria with dense connective tissue, no presenting gland, without separation between them (a), muscle wrap with bundles of muscles in vertical direction (b). In C, D, E, and F scanning electron microscopy (SEM) of the surface of the esophageal mucosa. In C cranial esophageal region, taste corpuscles to the center sensory cells with microvilli immersed in the central pore (white arrow head). In D and E presents the medial esophageal region, folds (db) and interdigitations (s) in different directions wrapped by cuboid-shaped epithelial cells (*), mucus release (arrow), and in F caudal portion of the esophageal surface is wrapped of epithelial cells with digitiform micro-saliences (md).

Morphologically, the acidophilic cells presented cubiform shape, enucleated, plasma membrane well demarcated and acidophylic cytoplasm of granulous aspect. The basophilic cells are larger, presenting round-shaped and basophilic cytoplasm. The epithelial cells present varied morphology from columnar to cuboidal (Fig. 2 B) and they are observed in greater number in places where the epithelium is composed of up to two layers of cells. Goblet cells (Fig. 2 A) are observed in smaller numbers, nucleated, presenting a light halo around the nucleus, being usually located in the medial region of the epithelium. The lamina propria of the mucosa and submucosa (Fig. 2 B) are composed of dense, not presenting gland, well-developed connective tissue and with no separation between them. Bundles of smooth muscles infiltrating the connective tissue of the mucous tunica were also observed, being more evident in the cranial portion of the esophagus. The muscle wrap is formed by bundles of muscles in a vertical direction.

By scanning electron microscopy (SEM) it was observed that esophageal mucosa is composed of numerous folds and sulcus in different directions (Fig. 2 D, E), and it is wrapped by epithelial cuboid-shaped cells, with the apical portion of some cells discontinued, suggesting the release of cytoplasmic content to lumen in the form of cellular explosion, and large amount of mucus on the surface and inside the interdigitations of the mucosa. However, in the cranial portion of the esophagus, mucosal folds are not observed, and the lining epithelium is formed by cells rounded to polyhedral, with a large amount of mucus. Furthermore, numerous projections in taste bud-shapes (Fig. 2 C) are observed on the epithelial surface, randomly distributed by the epithelium line. Such projections are intra-epithelial structures formed by grouping of support cells of the pavement type and in the center, numerous sensory cells with microvilli immersed in the central pore (white arrowhead). However, the surface is formed by epithelial cells with digitiform micro-saliences in the caudal portion of the esophagus.

The stomach mucosa of the *A. gigas* is covered with the simple columnar epithelium with the presence of numerous crypts and gastric folds, of morphology, height and in different numbers between the portions. In the cardiac portion (Fig. 3 A), the crypts have the end of a rounded-to-round aspect, with the presence of smaller secondary crypts among them. The mucosal lamina propria is formed by glandular loose connective tissue, which is observed in the base region of the crypts.

**Fig. 3.**
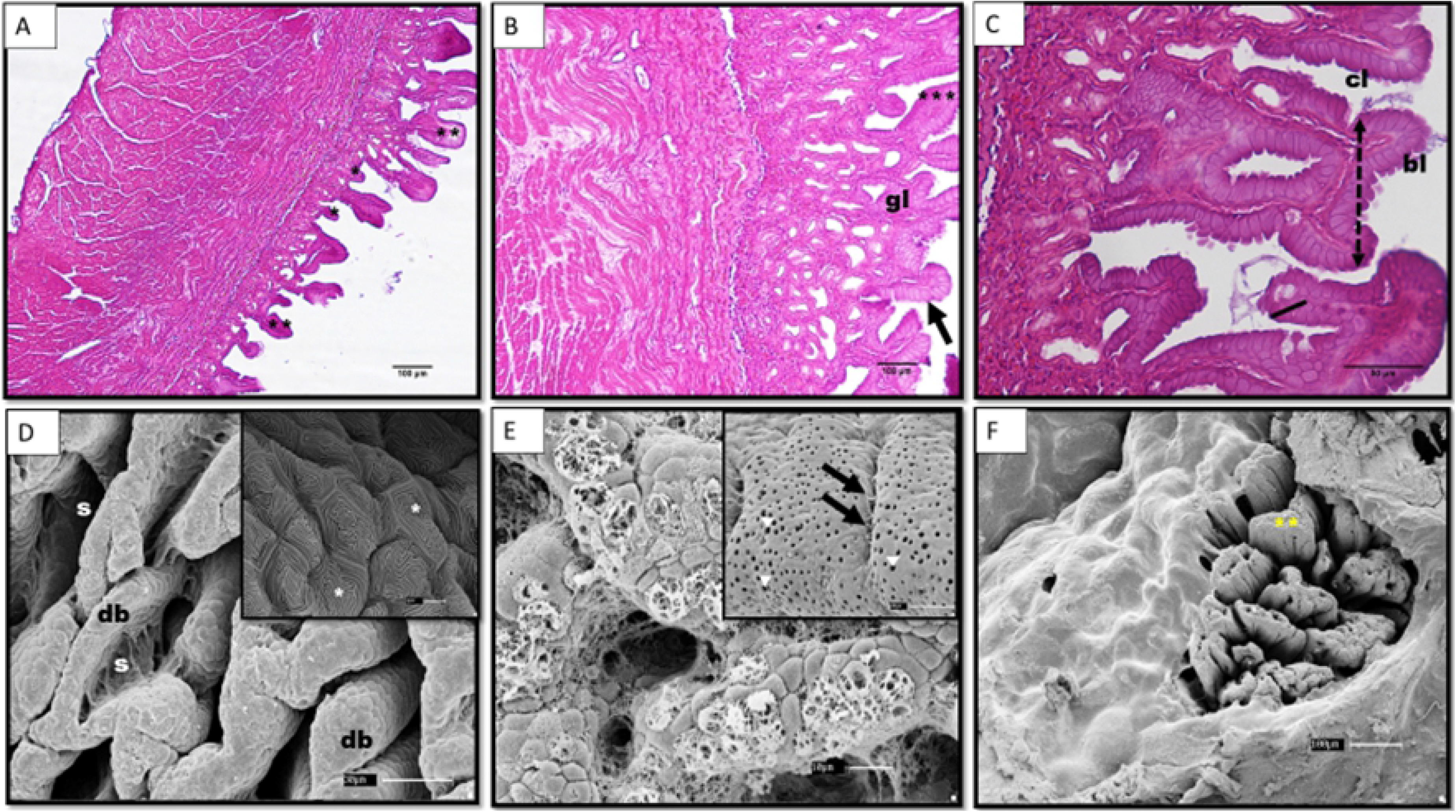
Photomicrography of *A. gigas* stomach. Cross-section in A, B, C (HE) in (ML). In A, cardiac portion, crypts with a rounded-to-round aspect (**), and secondary crypts (*), B in the fundus portion, column epithelial cells (arrow), and slightly eosinophilic cytoplasm with elevated and slender crypts and pointed extremity (***), in the mucosal lamina propria, more glands (gl) than connective tissue, these being larger than the cardiac portion. In C pyloric portion, baloon-shaped columnar epithelial cells (cl), epithelium hyperplasia (line), and thicker and shorter gastric crypts (dotted double arrow). In D, E, F in the SEM of the glandular surface of the stomach mucosa, in the cardiac portion (D) numerous folds (db) and interdigitations (s) and inside the interdigitations, cuboid epithelial cells replaced by epithelial cells with the digitiform micro-salience plasma membrane (*). E, cubiform epithelial cells with the presence of micro orifices (arrowheads), and at the ends of the mucosa folds the epithelial cells present dilation of the intercellular spaces, which allows the visualization of desmosome (arrows). In F, the pyloric portion presents flatter and thicker folds with presence of mucus (**).

In the fundus portion (Fig. 3 B), the epithelial cells of wrapping are columnar, with slightly eosinophilic cytoplasm and the crypts are elevated and slender with the pointed end, besides being more frequent compared to the cardiac portion. In the mucosal lamia propria, a greater number of glands than connective tissue is observed, which are larger than the cardiac portion.

In the pyloric portion the epithelial cells present a columnar to balloon form, with areas of epithelium hyperplasia, with up to three layers of cells. Such cells are also observed in glands located in the mucosal lamina propria. The gastric crypts of this region are thicker and shorter.

There is no separation between the mucous wrap and mucosal lamina propria, however the connective tissue of the mucous wrap is formed of thicker and denser collagen than the mucosal lamina propria. The muscular wrap is formed of bundles of smooth muscles arranged in two layers, horizontally and vertically. The ultramicroscopic scanning study of the surface of the paiche’s stomach mucosa presented it is formed of numerous folds and interdigitations (D). These are overlaid by cuboid epithelial cells (E) with micro-orifices (arrowhead) on the surface of the plasma membrane of these cells.

At the ends of the mucosa folds, epithelial cells present dilation of the intercellular spaces, which allows the visualization of desmosome (arrows). Inside the interdigitations, cuboid epithelial cells are replaced by epithelial cells with the plasma membrane (Fig. 3 D) with digitiform micro-salience. These cells reduce in quantity in the fundus portion and they were not found in the pyloric part. Mucus was observed in the three regions of the stomach; however the pyloric and cardiac portion were the sites with the highest amount, also presenting mucus and more flattened villi (Fig. 3 F).

## Discussion

The anatomical arrangement of the digestive system of the paiche (*A. gigas)* is similar to other teleosts. However, the stomach has varied shapes and regions; structurally adapted to food types. Anatomically, the stomach of the *A. gigas* is elongated in J-shape, similar to the species *Rhamdia quelen* [7], with a chemical region, and another mechanical, highly muscularized, with crushing function of the ingested food, also observed in piscivore stomachs [12].

The esophagus of the *A. Gigas* is a muscular organ, with several folds of mucosa and a fan-shaped dilation in the cranial portion with marked reduction of lumem in the medial portion, thus enabling distension during ingestion of large foods. Generally, this anatomical aspect has been observed in other carnivorous fish such as *Hoplias malabaricus* [13], *Salminus brasiliensis* [14], and *H. lacerdae* [15], as well as in other teleosts with different eating habits [15–17]. According to Rodrigues and Cargnin-Ferreira [10] and Scadeng et al. [18] the several amount of folds of mucosa supports the integrity of the esophageal wall, especially during the sudden distension motivated by quick ingestion.

By the scanning microscopy and light microscopy, a variation in the morphology of epithelial cells of esophageal wrapping was observed, thus, in the cranial portion, the mucosa was covered by rounded to polyhedral cells with the presence of taste corpuscle and in the other portions, secretory epithelial cells. The gustatory system of teleosts is activated by water-soluble substances, and these are involved in detection, selection, and ingestion.

According to Santos et al. [19] there is a close relationship between the distribution pattern of taste corpuscles and the way fish locate and select foods, in carnivores usually present in the buccopharyngeal cavity and little observed in the esophagus. The presence of such corpuscles in the cranial portion of the paiche esophagus, allow them to eject the already ingested food and even the eversion of the stomach when they feed on something strange or not palatable, as well as observed in salmon [20], and other teleosts [19].

In addition to the taste corpuscles, three different cell types were also observed in the stratified epithelium of the esophagus, mainly in the medial and caudal portions, the basophilic, acidophilic and claviform mucous cells. The epithelium of the teleosts is quite variable according to species, which may vary from stratified squamous to stratified columnar with three cell types: secretory cells of neutral, acidic and mixed mucopolysaccharides [21], such cells were also observed in the paiche’s esophagus of this study. According to Rodrigues and Cargnin-Ferreira [10] the goblet cells produce mucus that protect the esophageal wall against chemical and mechanical action during the passage of food. Nevertheless, acid mucus produced by acidophilic cells, is associated with binders capable of aggregating food particles, besides efficiently protecting the epithelium [22].

Bertin [23] describes on stomach of the teleosts different anatomical conformations and it can be grouped into three types: siphonal, cecum, and linear. The siphonal stomach has two branches, one descending, corresponding to the cardiac and the pyloric branch that ascends. The cecum stomach on the other hand, has two branches that interpose in the union region, the blind and rectilinear fundus and a muscular tube. However, it was possible to anatomically and histologically differentiate the stomach of the *Arapaima gigas* in three distinct portions: the cardiac, fundus, and pyloric, all glandular and presenting different histological characteristics, mainly at the ends of the gastric crypts, as well as the shape and number of glands of the mucosal lamina propria. Such morphological conformations corroborate the classification suggested by Cardoso et al. [24], that divide the stomach according to the anatomical structures and the presence or not of gastric glands in three parts, as described in the paiche.

The study of the digestive tract of several fish species, particularly teleosts, has attracted the attention of many researchers, mainly due to the wide structural variation with diversity of eating habits and behaviors, which are not found in other vertebrates. Based on anatomical and histological characteristics of the digestive tract, we can comprehend the feeding of fish [25]. The observation of gastrolithic pebbles inside the paiches’ stomach of this study may be associated with an adaptation to the type of pelletized food. For Magalhães et al. [26] the presence of pebbles in the stomach is related to the maceration process of pelletized foods, aided by stomach peristaltic waves. However, considering the low number of specimens in these studies, further research with native and captive fish are necessary to verify whether the presence of these pebbles is a food adaptation. On the ultramicroscopic scanning study of the surface of the paiche stomach mucosa, it was observed that it is formed by folds and interdigitations, also it is overlaid with different cell types such as cuboid epithelial cells, which secrete mucus, and epithelial cells with digitiform micro-salience. Cells with digitiform micro-salience were also observed in the caudal portion of the paiche’s esophagus. Morphological studies based on *Colossoma macropomum* also observed the presence of such cells, overlaying the esophagus [27]. According to Carrassón [28], these cells protect the epithelial surface and anchor the mucus secreted by goblet cells, besides being associated with fluids and ions absorption.

## Conclusion

Based on anatomical and morphological aspects observed in the esophagus of this study, the anatomical fan-shaped of the cranial portion, as well as the presence of taste corpuscles, enables the storage of food before passing to the stomach and the neurostimulation of flavors by taste corpuscles stimulate the bundles of muscle present in the submucosa wrap, carrying out the regurgitation or deglutition. Then morphology found was consistent with the eating habits and management of distinct characteristics of the digestive tract. However, these studies were carried out in juvenile fish, raised in aquaculture systems, fed with pelletized feed, which may not be observed in *A. gigas* in natural environment.

## Acknowledgments

We would like to thank CAPES/PROCAD Amazônia for the scholarship conceptions and for the resources made available for the study to be carried out.

## Author contributions

**Conceptualization:** Keila Silva Pinto, Luana Félix de Melo, Julia Bastos de Aquino, Jerônimo Vieira Dantas Filho, Maria Angelica Miglino, Sandro de Vargas Schons, Rose Eli Grassi Rici

**Data curation:** Keila Silva Pinto, Luana Félix de Melo, Julia Bastos de Aquino, Jerônimo Vieira Dantas Filho, Maria Angelica Miglino, Sandro de Vargas Schons, Rose Eli Grassi Rici

**Formal analysis:** Keila Silva Pinto, Luana Félix de Melo, Julia Bastos de Aquino, Jerônimo Vieira Dantas Filho, Maria Angelica Miglino, Sandro de Vargas Schons, Rose Eli Grassi Rici

**Funding acquisition:** Keila Silva Pinto, Luana Félix de Melo, Julia Bastos de Aquino, Jerônimo Vieira Dantas Filho, Maria Angelica Miglino, Sandro de Vargas Schons, Rose Eli Grassi Rici

**Investigation:** Keila Silva Pinto, Luana Félix de Melo, Julia Bastos de Aquino, Jerônimo Vieira Dantas Filho, Maria Angelica Miglino, Sandro de Vargas Schons, Rose Eli Grassi Rici

**Methodology:** Keila Silva Pinto, Luana Félix de Melo, Julia Bastos de Aquino, Jerônimo Vieira Dantas Filho, Maria Angelica Miglino, Sandro de Vargas Schons, Rose Eli Grassi Rici

**Project administration:** Luana Félix de Melo, Julia Bastos de Aquino, Maria Angelica Miglino, Sandro de Vargas Schons, Rose Eli Grassi Rici

**Resources:** Keila Silva Pinto, Luana Félix de Melo, Julia Bastos de Aquino, Jerônimo Vieira Dantas Filho, Maria Angelica Miglino, Sandro de Vargas Schons, Rose Eli Grassi Rici

**Supervision:** Sandro de Vargas Schons

**Validation:** Keila Silva Pinto, Luana Félix de Melo, Julia Bastos de Aquino, Jerônimo Vieira Dantas Filho, Maria Angelica Miglino, Sandro de Vargas Schons, Rose Eli Grassi Rici

**Visualization:** Keila Silva Pinto, Luana Félix de Melo, Julia Bastos de Aquino, Jerônimo Vieira Dantas Filho, Maria Angelica Miglino, Sandro de Vargas Schons, Rose Eli Grassi Rici

**Writing – original draft:** Keila Silva Pinto, Sandro de Vargas Schons

**Writing – review & editing:** Keila Silva Pinto, Luana Félix de Melo, Julia Bastos de Aquino, Jerônimo Vieira Dantas Filho, Maria Angelica Miglino, Sandro de Vargas Schons, Rose Eli Grassi Rici

## References

1. Amaral E, Sousa IS, Gonçalves ACT, Braga R, Ferraz P, Carvalho G. Manejo de pirarucus (*Arapaima gigas*) em Lagos de Várzea de Uso Exclusivo de Pescadores Urbanos e Ribeirinhos. Série Protocolos de Manejo de Recursos Naturais, Instituto de Desenvolvimento Sustentável Mamirauá – IDSM/OS/MCTI Programa de Manejo de Pesca (PMP). 2011.

2. Cavali J, Nunes CT, Dantas Filho, JV, Nóbrega BA, Pontuschka RB, Souza MLR, Porto MO. Chemical composition of commercial cuts of pirarucu (*Arapaima gigas*) processed in different weight classes in the Western Amazon. Revista Iberto-Americana de Ciências Ambientais. 2021; 12(4).

3. Nelson LS. Fishes of the world. 3. Ed. New York: John Wiley & Sons. 1994.

4. Nunes, ESCL, Franco RM, Mársico ET, Neves MS. Qualidade do pirarucu (*Arapaima gigas* Shing, 1822) salgado seco comercializado em mercados varejistas. Revista do Instituto Adolfo Lutz. 2012; 71: 520–529.

5. Azevedo PB, Morey GAM, Malta JCO. 2017. Mortalidade de juvenis de *Arapaima gigas* (Pisces: Arapaimidae) de piscicultura do norte do Brasil, causadas por *Hysterothylacium* sp. e *Goeziaspinulosa* (Nematoda: Anisakidae). Biota Amazônia. 2017; 7: 103–107.

6. Dantas Filho JV, Cavali J, Nunes CT, Nóbrega BA, Gasparini LRF, Souza MLR, Porto MO, Rosa BL, Gasparotto PHG, Pontuschka RB. Proximal composition, caloric value and price-nutrients correlation of comercial cuts of tambaqui (*Colossoma macropomum*) and pirarucu (*Arapaima gigas*) in diferente body weight classes (Amazon: Brazil). Research, Society and Development. 2021; 10: e23510111698.

7. Hernández DR, Pérez Gianeselli M, Domitrovic HA. Morphology, histology and histochemistry of the digestive system of South American Catfish (*Rhamdia quelen*). Internacional Journal of Morhology. 2009; 27: 105–111.

8. Løkka G, Austbø L, Falk K, Bjerkås I, Koppang EO. Intestinal morphology of the wild Atlantic salmon (*Salmo salar*). Journal Morphology. 2013; 274: 859–876.

9. Faccioli CK, Chedid RA, Amaral AC; Franceschini VB, Vicentini CA. Morphology and histochemistry of the digestivetract in carnivorous freshwater *Hemisorubim platyrhynchos* (Siluriformes: Pimelodidae). Micron. 2014; 64: 10–19.

10. Rodrigues APO, Cargnin-Ferreira E. Morphology and histology of the pirarucu (*Arapaima gigas*) digestive tract. Internacional Journal Morphology. 2017; 35: 950–957.

11. Buddington RK, Krogdahl A, Bakke-mckellep AM. The intestines of carnivorous fish: structure and functions and the relations with diet. Acta Physiologica Scandinavica. 1997; 638: 67–80.

12. Resende EK, Pereira RAC, Almeida VLL, Silva AG. Peixes onívoros da planície inundável do rio Miranda, Pantanal, Mato Grosso do Sul, Brasil. Corumbá: Boletim de Pesquisa Embrapa Pantanal. 2000; 60 p.

13. Menin E, Mimura OM. Anatomia comparativa do esôfago de seis peixes Teleostei de água doce de distintos hábitos alimentares. Revista Ceres. 1993; 15: 334–369.

14. Rodrigues SS, Menin E. Anatomia do tubo digestivo de *Salminus brasiliensis* (Cuvier, 1817) (Pisces, Characidae, Salmininae). Biotemas. 2008; 21: 65–75.

15. Hani YMI, Marchand A, Turies C, Kerambrum E, Palluel O, Bado-Nilles A, Beaudouin R, Porcher J, Geffard A, Dedourge-Geffard O. Digestive enzymes and gut morphometric parameters of threespine stickleback (*Gasterosteus aculeatus*): Influence of body size and temperature. PLOS ONE. 2018; 13: e0194932.

16. Schuingues CO, Lima MG, Lima AR, Martins DS, Costa GM. Anatomy of the bucopharyngeal cavity of *Sorubim trigonocephalus* (Siluriformes, Osteichthyes). Pesquisa Veterinária Brasileira. 2013; 33:1256–1262.

17. Fernandes MN, Cruz AL, Costa OTF, Perry SF. Morphometric partitioning of the respiratory surface area and diffusion capacity of the gills and swim bladder in juvenile Amazonian air-breathing fish, *Arapaima gigas*. Micron. 2012; 43: 961–970.

18. Scadeng M, McKenzie C, He W, Bartsch H, Dubowitz DJ, Stec D, Leger JS. Morphology of the Amazonian Teleost Genus Arapaima Using Advanced 3D Imaging. Frontiers in Physiology. 2020; 11.

19. Santos AFGN, Santos LNS, Araújo FG. Digestive tract morphology of the Neotropical piscivorous fish *Cichla kelberi* (Perciformes: Cichlidae) introduced into an oligotrophic Brazilian reservoir. Biologia Tropical. 2011; 59: 1245–1255.

20. Gioda CR, Pretto A, Freitas CS, Leitemperger J, Loro VL, Lazzari R, Lissner LA, Baldisserotto B, Salbego J. Different feeding habits influence the activity of digestive enzymes in freshwater fish. Ciência Rural. 2017; 47: e20160113.

21. Santo ML, Arantes FP, Pessali TC, Santos JE. Morphological, histological and histochemical analysis of the digestive tract **of** Trachelyopterusstriatulus (Siluriformes: Auchenipteridae). Zoologia. 2015; 32: 296–305.

22. Morrison CM et al. 1987. Histology of the Atlantic cod, *Gadus morhua*: an atlas. Pt. 4: Eleutheroembryo and larva.

23. Bertin L. Appareil digestif. In: Grassé PP. Traite de zoologia. 1958; 13: 1248–1302.

24. Cardoso NN, Firmiano EMS, Gomes ID, Nascimento AA, Sales A, Araújo FG. Histochemical and immunohistochemical study on endocrine cells (5HT, GAS, and SST) of the gastrointestinal tract of a teleost, the characin *Astyanax bimaculatus*. Acta Histochemica. 2015; 117: 595–604.

25. Machado MRF, Souza HO, Souza VL, Azevedo A, Goitein R, Nobre AD. Morphological and anatomical characterization of the digestive tract of *Centropomus parallelus* and *C. Undecimalis*. Acta Scientiarum Biological Sciences. 2013; 35: 467–474, 2013.

26. Magalhães Junior FO, Santos MJM, Allaman IB, Soares Junior IJ, Silva RF, Braga LGT. Digestible Protein Requirement of Pirarucu Juveniles (*Arapaima gigas*) Reared in Outdoor Aquaculture. Journal of Agricultural Science. 2017; 9(9).

27. Cavali, J, Dantas Filho JV, Nóbrega BA, Andrade LHV, Pontuschka RB, Gasparotto PHG, Francisco RS, Campeiro Junior LD, Porto MO. Benefits of adding virginiamycin to *Arapaima gigas* (Schinz, 1822) diet cultivated in the Brazilian Amazon. Scientifica. 2020; doi: https://doi.org/10.1155/2020/5953720

28. Carrassón M, Grau A, Dopazo LR, Crespo S. A histological, histochemical and ultrastructural study of the digestive tract of Dentex dentex (Pisces, Sparidae). Histology and Histopathology. 2006; 21: 579–593

